# Vitamin D-binding protein is required for the maintenance of α-cell function and glucagon secretion

**DOI:** 10.1101/2019.12.19.881185

**Authors:** Katrina Viloria, Daniela Nasteska, Linford J.B. Briant, Silke Heising, Dean Larner, Nicholas H.F. Fine, Fiona B. Ashford, Gabriela da Silva Xavier, Maria Jiménez Ramos, Jocelyn E. Manning Fox, Patrick E. MacDonald, Ildem Akerman, Gareth G. Lavery, Christine Flaxman, Noel G. Morgan, Sarah J. Richardson, Martin Hewison, David J. Hodson

## Abstract

Vitamin D-binding protein (DBP) or GC-globulin carries vitamin D metabolites from the circulation to target tissues. DBP expression is highly-localized to the liver and pancreatic α-cells. While DBP serum levels, gene polymorphisms and autoantigens have all been associated with diabetes risk, the underlying mechanisms remain unknown. Here, we show that DBP regulates α-cell morphology, α-cell function and glucagon secretion. Deletion of DBP led to smaller and hyperplastic α-cells, altered Na^+^ channel conductance, impaired α-cell activation by low glucose, and reduced rates of glucagon secretion. Mechanistically, this involved reversible changes in islet microfilament abundance and density, as well as changes in glucagon granule distribution. Defects were also seen in β-cell and δ-cell function. Immunostaining of human pancreata revealed generalized loss of DBP expression as a feature of late-onset and longstanding, but not early-onset type 1 diabetes. Thus, DBP is a critical regulator of α-cell phenotype, with implications for diabetes pathogenesis.

**HIGHLIGHTS:**

- DBP expression is highly-localized to mouse and human α-cells
- Loss of DBP increases α-cell number, but decreases α-cell size
- α-cells in DBP knockout islets are dysfunctional and secrete less glucagon
- DBP expression is decreased in α-cells of donors with late-onset or longstanding type 1 diabetes

## INTRODUCTION

Vitamin D-binding protein (DBP), a 52–59 kDa protein also known as group-specific component of serum (GC-globulin), is the primary plasma carrier for circulating vitamin D and its metabolites (White and Cooke, 2000). Expression of *GC/Gc*, which encodes DBP, is highly-expressed in the liver of all mammals (Feldman et al., 2017), in keeping with the function of this organ to convert sterol-derivatives such as cholecalciferol (Vitamin D_3_) into pre-hormone 25-OH vitamin D (25(OH)D) (Bikle, 2014). DBP is also expressed in the pancreatic islets. Recent studies have shown that *GC/Gc* is highly-expressed in purified mouse and human α-cells (Ackermann et al., 2016; Adriaenssens et al., 2016; Qiu et al., 2017; Segerstolpe et al., 2016), and is upregulated in de-differentiated β-cells (Kuo et al., 2019). Since the *GC* promoter region contains cell type-selective open chromatin regions, *GC* can be classified as an α-cell signature gene, similarly to prototypical hits such as *ARX, GCG, IRX2* and *DPP4* (Ackermann et al., 2016; Lam et al., 2019). Despite these findings, the role of DBP in the regulation of glucagon release, and in wider aspects of islet function, remains enigmatic.

Evidence that the effects of DBP in α-cells are unrelated to serum vitamin D transport comes from studies in vitamin D-deficient patients who show no improvement in insulin-induced glucagon output upon vitamin D repletion (Gedik and Akalin, 1986). Moreover, a patient harboring a rare mutation in *GC* showed no symptoms of vitamin D deficiency, despite low plasma levels of 25(OH)D, arguing that it is the free form of 25(OH)D that dictates many of the non-classical actions of vitamin D (Chun et al., 2014; Henderson et al., 2019). Alongside its role in 25(OH)D transport, DBP is also a major actin scavenger (Harper et al., 1987). Following disassembly of polymerized actin by gelsolin, DBP traps monomeric filaments using its three domains as a clamp (Otterbein et al., 2002). Pertinently, ephrin-A forward signaling has been shown to inhibit glucagon secretion through increases in F-actin density (Hutchens and Piston, 2015), and the appearance of regulated glucagon secretion in re-aggregated islets coincides with normalization of F-actin levels (Reissaus and Piston, 2017).

Linking DBP with type 2 diabetes (T2D) risk, *GC* variants are associated with elevated fasting glucose, fasting insulin levels and responses to oral glucose tolerance (Baier et al., 1998; Hirai et al., 2000; Iyengar et al., 1989; Szathmary, 1987). Results, however, tend to be conflicting, likely reflecting heterogeneity introduced by ethnicity and environment (Malik et al., 2013; Wang et al., 2014). The concept that DBP might also be involved in type 1 diabetes (T1D) risk is supported by retrospective cross-sectional analysis of 472 individuals showing that serum DBP levels were lowest in patients with T1D (Blanton et al., 2011). Using gene expression-based genome-wide association studies, DBP was subsequently identified as a novel T1D autoantigen (Kodama et al., 2016). The same authors showed that T-cell reactivity against DBP was increased in NOD mice, and that humans with T1D possess specific DBP autoantibodies (Kodama et al., 2016). Together, these studies suggest that DBP is likely to be associated with altered diabetes risk in humans.

Here, we sought to establish the role of DBP in α-cell phenotype, function and diabetes risk by combining studies in knockout mice with staining of pancreata from T1D donors and age-matched controls. We show that DBP is critical for proper α-cell function and glucagon secretion, with related effects for δ-cell morphology and insulin release. We further show that glucagon and DBP expression decrease in α-cells of individuals with late-onset or longstanding T1D, but not in children with early-onset disease. As such, DBP should be considered as an essential component of the α-cell and the wider islet functional machinery with relevance for glucagon secretion during diabetes.

## RESULTS

### DBP is deleted in α-cells of DBP^-/-^ mice

Confocal imaging showed an intense DBP signal localized predominantly to GLU+ cells at the islet periphery in mice (Figure 1A). While DBP expression was clearly decreased in DBP^-/-^ animals, a faint signal could still be detected in some α-cells using fluorescent immunohistochemistry (Figure 1A). This likely reflects the sensitivity of the staining protocol used since non-fluorescent immunohistochemistry implied complete loss of DBP in DBP^-/-^ islets (Figure 1B). We wondered whether DBP was also expressed in other islet cell types, but might be obscured by the strong staining detected in α-cells. Therefore, immunostaining was repeated using a higher antibody concentration combined with more sensitive imaging settings to oversaturate signal in α-cells but not in other cells. Using this approach, weak DBP expression could be detected in the β-cell compartment, which was absent in islets from DBP^-/-^ mice (Figure 1C).

**Figure 1:**
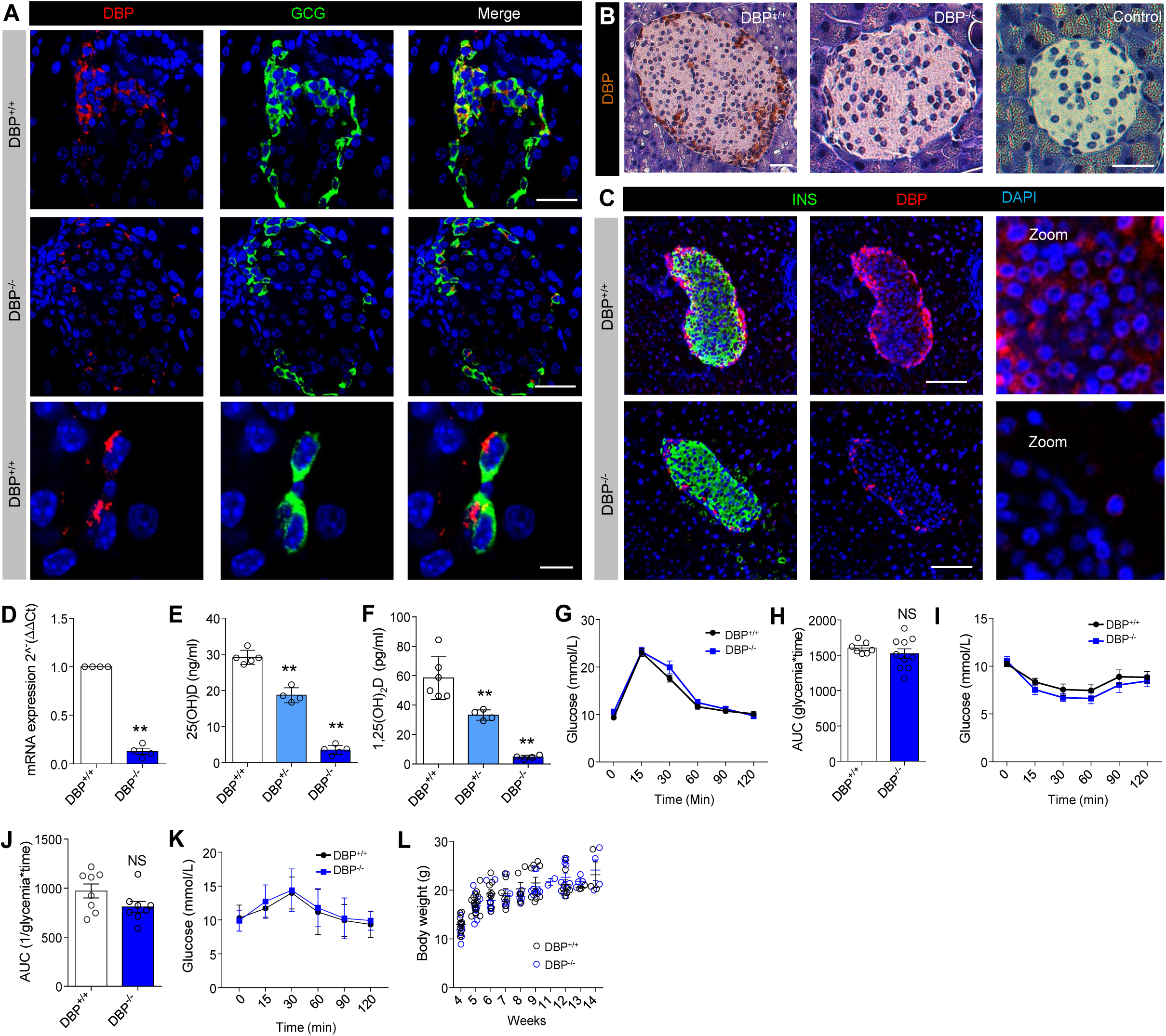
Phenotypic assessment of DBP^-/-^ mice. (A) Representative fluorescence immunohistochemistry image showing localization of DBP to α-cells and its specific loss in DBP^-/-^ animals (scale bar = 34 µm, top and middle panels) (scale bar = 12 µm, bottom panel) (n = 3-4 animals). (B) As for (A), but non-fluorescent DAB staining in DBP^+/+^ and DBP^-/-^ pancreatic sections (scale bar = 40 µm) (n = 2-3 animals). (C) Fluorescence immunohistochemistry showing DBP staining of β-cells, which is absent in pancreatic sections from DBP^-/-^ animals. Due to the relative strength of DBP expression in α-cells, the images have been overexposed to allow visualization of DBP in the non α-cell compartment (representative images are shown) (scale bar = 85 µm) (n = 13 islets, 3 animals). (D) Expression of *Gc*, which encodes DBP, is barely detectable in DBP^-/-^ islets using Taqman assays (n = 4-5 animals). (E and F) Serum 25(OH)D (D) and 1,25(OH)2D (E) levels are ∼2-fold and 4-fold lowered in DBP^+/-^ and DBP-/-animals, respectively (n = 4-6 animals) (one-way ANOVA with Bonferroni’s multiple comparisons test). (G and H) Glucose tolerance curves (G) and AUC (H) are similar in DBP^+/+^ and DBP^-/-^ mice (n = 7-11 animals) (two-way repeated measure ANOVA with Bonferroni’s multiple comparisons test, or unpaired t-test). (I and J) Insulin sensitivity is similar in DBP^+/+^ and DBP^-/-^ mice (I), as also shown by the AUC (J) (n = 11 animals) (two-way repeated measure ANOVA with Bonferroni’s multiple comparisons test, or unpaired t-test). (K and L) Pyruvate tolerance (K) and body weight/growth curve (L) is similar in DBP^+/+^ and DBP^-/-^ mice (n = 3-4) (two-way repeated measure ANOVA or unpaired t-test). Line graphs show mean ± SEM. Bar graphs show scatter plot with mean ± SEM. *P<0.05, **P<0.01 and NS, non-significant. DBP, Vitamin D-binding protein; DAB, 3,3′-Diaminobenzidine; GLU, glucagon; AUC, area-under-the-curve (AUC).

*Gc* was undetectable using specific Taqman assays (Figure 1D), and circulating 25(OH)D and hormonal 1,25-(OH)_2_ vitamin D (1,25(OH)_2_D) levels were ∼50% decreased in heterozygous DBP^+/-^ mice, and virtually undetectable in homozygous DBP^-/-^ littermates (Figure 1E and 1F). Despite low levels of 25(OH)D and 1,25(OH)_2_D, DBP^-/-^ animals do not show signs of vitamin D deficiency unless placed on a vitamin D-deficient diet (Safadi et al., 1999).

Metabolic phenotyping of DBP^-/-^ mice at 8-12 weeks revealed normal glucose tolerance (Figure 1G and H), insulin sensitivity (Figure 1I and 1J) and pyruvate tolerance versus littermate controls (Figure 1K). Growth curves and adult body weight were similar between genotypes (Figure 1L). Thus DBP^-/-^ mice possess an apparently normal metabolic phenotype, without changes in insulin sensitivity or hepatic gluconeogenesis that could non-autonomously influence α-cell mass/function. With this in mind, we proceeded to conduct the remainder of the studies in isolated pancreata and islets.

### Deletion of DBP leads to abnormal islet morphology

Cell resolution immunostaining of entire pancreatic sections showed no changes in α-cell or β-cell mass following loss of DBP (Figure 2A and B). While detailed morphometric analyses of individual DBP^-/-^ islets revealed an increase in α-cell number (Figure 2C and D) (also apparent in Figure 1A), this was accompanied by a decrease in α-cell size (Figure 2E), maintaining the area occupied by α-cells (Figure 2F). Although a small but significant increase in β-cell number was apparent (Figure 2G), no changes in β-cell size (Figure 2H) or area occupied by β-cells (Figure 2I) were detected between DBP^-/-^ and DBP^+/+^ animals. However, a ∼2-fold decrease in δ-cell mass was detected (Figure 2J and 2K), along with a reduction in the size of individual δ-cells (Figure 2L and 2M).

**Figure 2:**
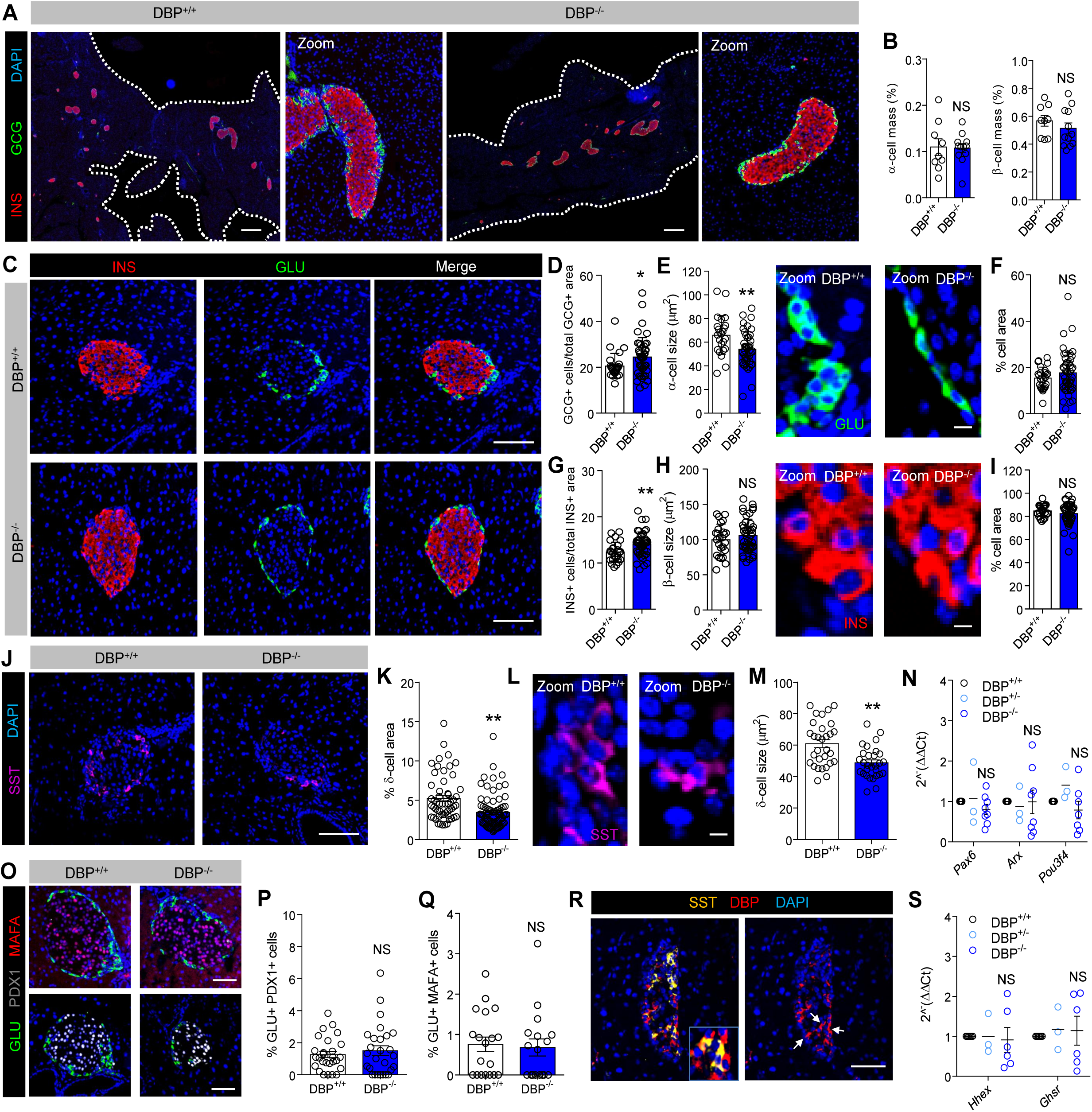
DBP alters α-cell and δ-cell number and size. (A and B) Cell resolution reconstruction of pancreatic sections reveals no differences in α-cell and β-cell mass in DBP^+/+^ and DBP^-/-^ mice (A), quantified in bar graph (B) (scale bar = 530 µm) (representative images are shown; inset is a zoom showing maintenance of cellular resolution in a single image) (n = 9-12 sections from 3-4 animals) (unpaired t-test). (C-I) Morphological analyses of DBP^-/-^ islets (C) reveal increased α-cell number (D), decreased α-cell size (E, representative images shown in right panel) but normal area occupied (F). By contrast, β-cell number is increased (G), although β-cell size (H, representative images shown in right panel) and area (I) are unchanged (scale bar in D = 85 µm; scale bar in E and H = 10 µm) (n = 24-45 islets from 3-4 animals) (panel E and H are zooms of panel C to better show α-cell and β-cell size) (DAPI is shown in blue) (unpaired t-test). (J-M) δ-cell proportion (J and K) (n = 55–79 islets from 4-5 animals) and size (L and M) (n = 29-30 islets from 3 animals) are decreased in DBP^-/-^ islets (scale bar in J = 85 µm; scale bar in L = 10 µm) (representative images are shown; panel L is a zoom of panel J to better show δ-cell cell size) (unpaired t-test). (N) Expression levels of the α-cell differentiation markers *Arx, Pax6* and *Pou3f4* are similar in DBP^+/-^ and DBP^-/-^ islets (n = 3-6 animals) (one-way ANOVA with Bonferroni’s multiple comparisons test). (O-Q) No changes in the proportion of α-cells expressing PDX1 (O and P) or MAFA (O and Q) are detected in pancreatic sections from DBP^-/-^ versus DBP^+/+^ islets (scale bar = 85 µm) (representative images are shown) (n = 17-27 islets from 3 animals) (unpaired t-test). (R) DBP is expressed in a subpopulation of δ-cells (arrows show SST+/DBP+ cells) (n = 3 animals) (scale bar = 85 µm). (S) The δ-cell differentiation markers *Hhex* and *Ghsr* are not significantly downregulated in DBP^-/-^ islets (n = 3-6 animals) (one-way ANOVA with Bonferroni’s multiple comparisons test). Bar graphs show scatter plot with mean ± SEM. *P<0.05, **P<0.01 and NS, non-significant. DBP, Vitamin D-binding protein; GLU, glucagon; NS, insulin; SST, somatostatin.

Suggesting that loss of DBP is not associated with α-cell de-differentiation, mRNA levels for *Pax6, Arx* and *Pou3f4* was similar in islets from DBP^+/+^ and DBP^-/-^ mice (Figure 2N). No differences in the number of (very rare) cells co-staining for GLU/PDX1 or GLU/MAFA were detected (Figure 2O-Q), implying that adoption of a β-cell- or δ-cell-like fate by α-cells was unlikely. Moreover, a similar number of PCNA+ or proliferative α-cells was detected in DBP^-/-^ and DBP^+/+^ animals, suggesting normal cell turnover rates (Figure S1). While DBP was expressed in some but not all δ-cells (Figure 2R), expression of the δ-cell differentiation markers *Hhex* and *Ghsr* was unchanged in DBP^-/-^ islets (Figure 2S).

Thus, DBP^-/-^ islets are morphologically abnormal, containing smaller and more abundant α-cells alongside a modest contraction of the δ-cell compartment.

### DBP is critical for α-cell, δ-cell and β-cell function

Multicellular Ca^2+^ imaging of DBP^-/-^ islets showed large impairments in the activity of α-cells, identified by their responses to low glucose and epinephrine (Figure 3A-C) (Tian et al., 2011). This presented as a loss of α-cell activation by low glucose (Figure 3A), although some α-cells that remained active displayed characteristic Ca^2+^-spiking responses of elevated amplitude (Figure 3B and 3C). We also examined the ability of α-cells to coordinate Ca^2+^ responses within clusters throughout the imaged plane. Unexpectedly, α-cell population activity in DBP^-/-^ mice was more stochastic, with an almost 2-fold decrease in coordinated α-cell-α-cell activity (Figure 3D). This was not dependent on loss of a specific subset of α-cells, since topology of the communicating α-cells was unaffected by DBP loss (Figure 3E).

**Figure 3:**
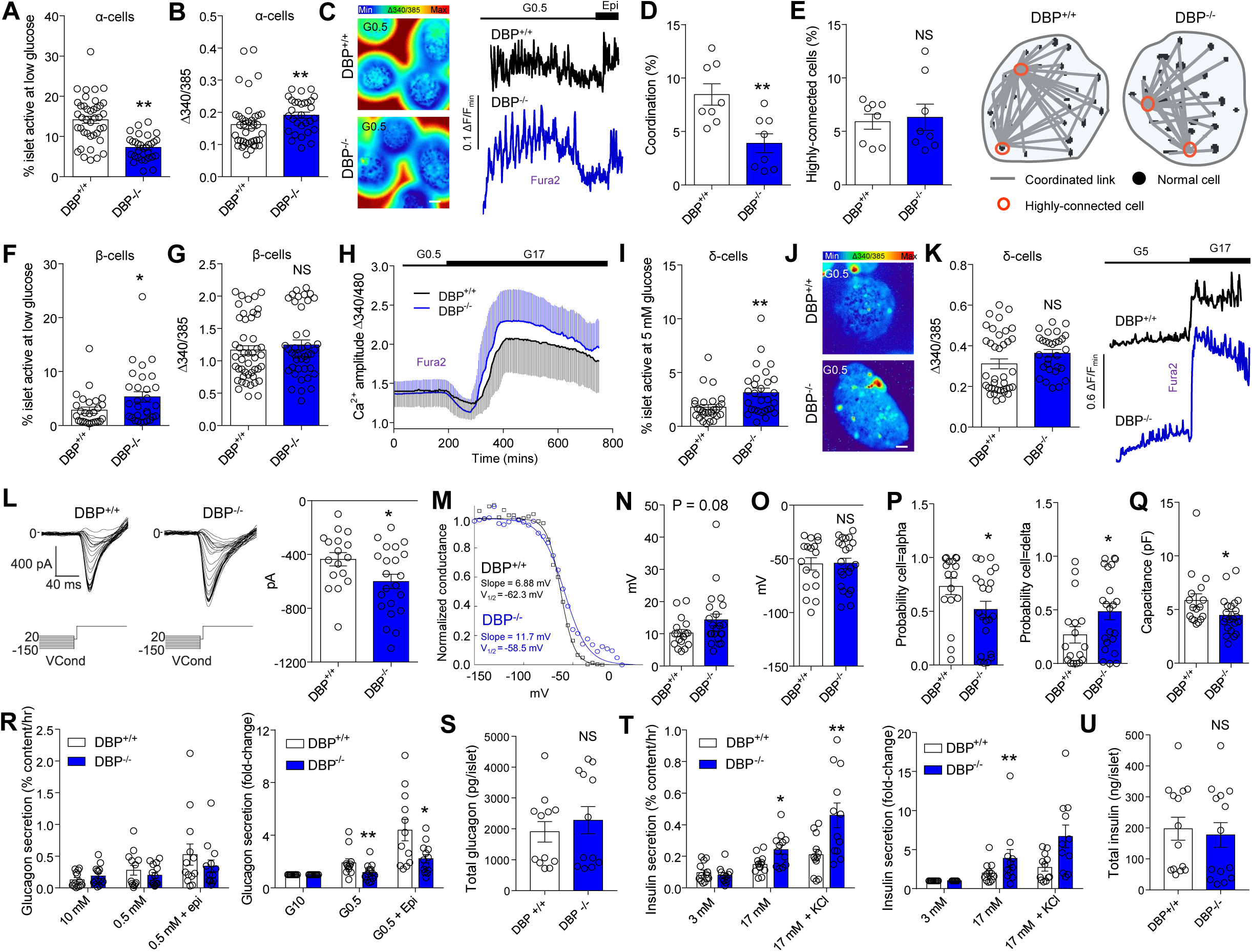
Dysregulated α-cell, β-cell and δ-cell function in islets lacking DBP. (A and B) Proportion of α-cells active at low (0.5 mM) glucose is decreased in DBP^-/-^ islets (A), although Ca^2+^ amplitude is increased in responsive cells (B) (n = 46-30 islets, 4 animals) (Mann-Whitney test). (C) Representative images (left) and traces (right) showing loss of α-cell activation in DBP^-/-^ islets (scale bar = 40 µm) (n = 46-30 islets, 4 animals). (D) Coordinated α-cell responses to low glucose are impaired in DBP^-/-^ islets (white box shows an episode of coordinated activity) (n = 8 islets, 4 animals) (unpaired t-test). (E) Topology of coordinated α-cells is similar in DBP^-/-^ versus DBP^+/+^ islets (n = 8 islets, 4 animals) (unpaired t-test). (F-H) Proportion of β-cells active at low (0.5 mM) (F) glucose is increased in DBP^-/-^ islets, despite intact responses to high (17 mM) glucose (G and H) (n = 27-29 islets, 4 animals) (unpaired t-test). (I-K) More δ-cells are active at 5 mM glucose in DBP^-/-^ compared to DBP^+/+^ islets (I and J), mounting Ca^2+^ spikes with a tendency toward increased amplitude (K) (representative Ca^2+^ images and traces are shown) (scale bar = 40 µm) (n = 28-29 islets, 4 animals) (unpaired t-test). (L) Representative patch-clamp recordings of α-cells (left) showing increased Na^+^ conductance in DBP^+/+^ islets (right) (n = 17-22 cells, 3 animals) (unpaired t-test). (M-O) Sigmoid plots of raw current data showing calculation of slope factor and half-maximal voltage (V_1/2_) for two cells (M). Summary data show a tendency toward increased slope factor for Na^+^ channel inactivation (N), but unchanged half maximal voltage (O) in DBP^-/-^ versus to DBP^+/+^ α-cells (n = 17-22 cells, 3 animals) (Mann-Whitney test). (P) Electrophysiological fingerprinting reveals decreased and increased probability of cells resembling an α-cell or δ-cell, respectively, in DBP^-/-^ islets (n = 17-22 cells, 3 animals) (Mann-Whitney test). (Q) α-cell capacitance is significantly reduced in DBP^-/-^ islets (n = 17-22 cells, 3 animals) (unpaired t-test). (R) Glucagon secretion is impaired in DBP^-/-^ islets in response to low (0.5 mM) glucose and low (0.5 mM) glucose + epinephrine (data are shown normalized to content or as fold-change) (n = 13-14 replicates, 10 animals) (two-way ANOVA with Bonferroni’s multiple comparisons test, or unpaired t-test). (S) Glucagon content is similar in DBP^+/+^ and DBP^-/-^ islets (n = 12 replicates, 8 animals) (unpaired t-test). (T) Insulin secretion in response to high (17 mM) glucose or high (17 mM) glucose + KCl is increased in DBP^-/-^ islets (data are shown normalized to content or as fold-change) (n = 12-13 replicates, 3 animals) (two-way ANOVA with ANOVA with Bonferroni’s multiple comparisons test, or unpaired t-test). (U) Insulin content is similar in DBP^+/+^ and DBP^-/-^ islets (n = 14 replicates, 3 animals) (unpaired t-test). Bar graphs show scatter plot with mean ± SEM. *P<0.05, **P<0.01 and NS, non-significant. DBP, Vitamin D-binding protein.

The same islets were also examined for changes in β-cell activity at both low (0.5 mM) and high (17 mM) glucose. A large increase in the proportion of β-cells active at low glucose was observed (Figure 3F), identified on the basis of their responsiveness to subsequent challenge with high glucose. However, no differences in β-cell activity were detected at high glucose (Figure 3G and 3H), suggesting the presence of intact glucose metabolism.

To record δ-cell activity, islets were imaged at 5 mM glucose before increasing concentration of the sugar to 17 mM. δ-cells were identified by their characteristic, rare, Ca^2+^-spiking patterns at 5 mM glucose (Figure 3K), which did not change in the presence of high glucose (compared to β-cells) (Shuai et al., 2016; Vierra et al., 2018). Suggesting the presence of abnormal function, the proportion of active δ-cells was increased in DBP^-/-^ islets (Figure 3I and 3J), possibly reflecting compensation for reduced δ-cell number. However, δ-cells that did respond displayed Ca^2+^ spikes of normal amplitude (Figure 3K).

### DBP is required for normal α-cell Na^+^ channel conductance

α–cells generate electrical activity, with Na^+^ channel inactivation properties playing an important role in determining glucagon secretion (Zhang et al., 2013). Using patch clamp electrophysiology, we therefore explored whether DBP influences α–cell Na^+^ channel function. As expected from the increased Ca^2+^ amplitude detected in these cells, Na^+^ currents were increased in α–cells lacking DBP (Figure 3L). However, the slope factor of Na^+^ inactivation also tended to be increased (Figure 3M and N), despite a similar half maximal voltage (Figure 3O), suggesting the presence of impairments in glucose-dependent α–cell activity.

We further subjected patch-clamp recordings from DBP^+/+^ and DBP^-/-^ islets to a mathematical model (Briant et al., 2017b). This model takes as input the electrophysiological data of a cell, and outputs a probability that the cell is an α–cell. While the model predicted α–cell phenotype in DBP^+/+^ islets with high probability, confidence was lower in recordings from DBP^-/-^ islets with the output favoring a more δ-cell-like (probability = 0.27 versus 0.48, DBP^+/+^ versus DBP^-/-^, respectively, P<0.05), but not β-cell-like (probability = 4.43 x 10^−7^ versus 1.15 x 10^−8^, DBP^+/+^ versus DBP^-/-^, respectively, non-significant) profile (Figure 3P).

Thus, α–cells lose their “electrophysiological identity”, become less α–cell-like, and resemble δ-cells following loss of DBP. Confirming the decrease in α–cell size detected using immunohistochemistry, membrane capacitance was significantly lower in cells predicted to be α–cells in DBP^-/-^ islets (Figure 3Q).

### DBP regulates glucagon and insulin secretion

DBP^+/+^ islets responded to low (0.5mM) glucose with a 2-fold increase in glucagon secretion (Figure 3R). By contrast, DBP^-/-^ islets showed a tendency toward increased basal glucagon levels and loss of glucagon secretion in response to low glucose (Figure 3R). Glucagon secretory responses to epinephrine were also significantly impaired in DBP^-/-^ islets, pointing toward a defect in exocytosis (Figure 3R). Suggesting the presence of normal glucagon biosynthesis, total levels of the hormone were similar between DBP^+/+^ and DBP^-/-^ littermates (Figure 3S).

Conversely to glucagon, glucose-stimulated insulin secretion was significantly increased in islets from DBP^-/-^ animals (Figure 3T). This effect was likely associated with improved exocytosis of the readily-releasable pool of insulin granules, since KCl-stimulated insulin secretion was similarly increased in DBP^-/-^ *versus* DBP^+/+^ islets (Figure 3T), despite equivalent insulin content (Figure 3U).

Thus, DBP is required for normal glucose-regulated glucagon secretion, and limits insulin secretion under high glucose stimulation.

### DBP mediates α-cell function through F-actin binding

DBP is a major actin scavenger and might exert effects on α-cell size and glucagon secretion by trapping monomeric actin (G-actin), which is needed to form polymerized actin (F-actin) (Dominguez and Holmes, 2011). To investigate DBP-actin interactions, high resolution snapshots were taken of islets stained with either phalloidin or DNAaseI to demarcate F-actin and G-actin, respectively. While F-actin density was unchanged in DBP^-/-^ islets, increases in intensity and fiber thickness were seen throughout the islet (Figure 4A-C), rather than restricted solely to α-cells. Conversely, G-actin levels tended to be reduced in DBP^-/-^ mice, again being evident throughout the islet (Figure 4D and E).

**Figure 4:**
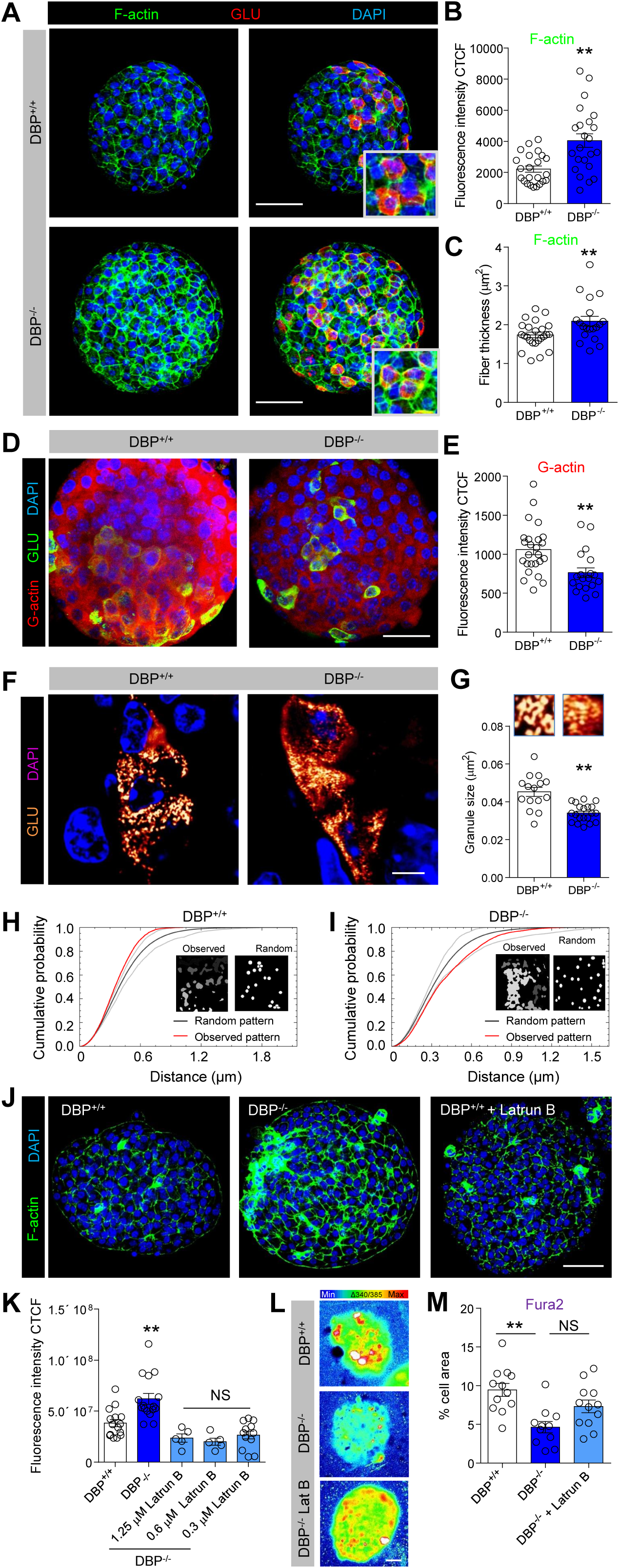
DBP scavenges actin in the islet and maintains glucagon granule morphology. (A-C) F-actin expression is increased following loss of DBP (A), quantified using fluorescence intensity (B) and fiber thickness (C) (representative images are shown) (scale bar = 53 µm) (n = 23 islets, 3 animals) (unpaired t-test). (D and E) As for (A-C), but representative images (D) and summary bar graph (E) showing decreased G-actin monomer expression (scale bar = 34 µm) (n = 19-25 islets, 5 animals) (unpaired t-test). (F and G) Representative super-resolution (∼ 140 nm lateral resolution) snapshots of glucagon granules (F), showing a 20% decrease in size (G) (magnified images from (F) are shown above each bar) (scale bar = 6 µm) (n = 13-15 islets, 3-4 animals) (unpaired t-test). (H and I) G-function analysis on actual and simulated glucagon granule distribution showing that glucagon granules tend to more diffusely scattered in DBP^-/-^ islets (actual and simulated distribution is inset). (J and K) Representative images (J) and bar graph (K) showing that F-actin levels in DBP^-/-^ islets can be restored to DBP^+/+^ levels using 0.3-1.25 µM Latrunculin B (scale bar = 53 µm) (n = 5-16 islets, 1-2 animals) (one-way ANOVA wih Dunnett’s post hoc test). (L and M) Representative images (L) and summary bar graph (M) showing that 0.3 µM Latrunculin B rescues α-cell responses to low glucose in DBP^-/-^ islets (scale bar = 25 µm) (n = 11-12 islets, 3-4 animals). Bar graphs show scatter plot with mean ± SEM. *P<0.05, **P<0.01 and NS, non-significant. DBP, Vitamin D-binding protein; GLU, glucagon; INS, insulin; Latrun B, Latrunculin B.

Since F-actin/G-actin ratio is important for regulated secretion (Kalwat and Thurmond, 2013), glucagon granule size and distribution were mapped in pancreatic slices using super-resolution imaging. Analysis of individual granules revealed a small but significant decrease in granule size in DBP^-/-^ mice, although area occupied was unchanged, pointing to an increase in granule number (Figure 4F and G). While glucagon granules tended to be clustered in DBP^+/+^ α-cells, they were more diffusely scattered throughout the cytoplasm in DBP^-/-^ tissue (Figure 4H and I).

Lastly, we investigated whether normal activity could be restored in DBP^-/-^ islets by reducing F-actin levels to DBP^+/+^ levels. Experiments were performed in the absence or presence of Latrunculin B, which prevents F-actin polymerization (Spector et al., 1983). A concentration-response showed that Latrunculin B was able to modulate F-actin within the range detected in DBP^+/+^ islets (Figure 4J). Notably, using 300 nM Latrunculin B to reduce F-actin in DBP^-/-^ islets down to DBP^+/+^ levels (Figure 4K), we were able to restore α-cell Ca^2+^ responses to low glucose (Figure 4L and M).

Together, these results suggest that DBP increases availability of monomeric G-actin, which is then able to polymerize to form F-actin, ultimately altering granule distribution and size, as well as α-cell function.

### DBP and glucagon expression are decreased in late-onset and longstanding T1D donors

Islet α-cells persist in T1D, but display reduced glucose responsiveness (Brissova et al., 2018; Gerich et al., 1973), which could be associated with changes in DBP expression. We therefore examined whether DBP levels changed in T1D, initially using pancreatic sections from the Exeter Archival Diabetes Biobank. Immunohistochemistry was performed on sections from donors with early- and late-onset T1D, together with their age-matched controls. In donors without diabetes, DBP was highly localized to α-cells (Figure 5A), as previously shown (Kodama et al., 2016; Lam et al., 2019), although we were also able to detect very faint expression in β-cells, as for mouse (Figure S3). In sections from donors with T1D, a similar pattern of DBP immunostaining was observed (Figure 5B). While a small but significant decrease in glucagon expression was seen in islets of early-onset T1D donors (Figure 5C), this was not accompanied by changes in DBP staining (Figure 5D) or proportion of α-cells immunopositive for DBP (Figure 5E).

**Figure 5:**
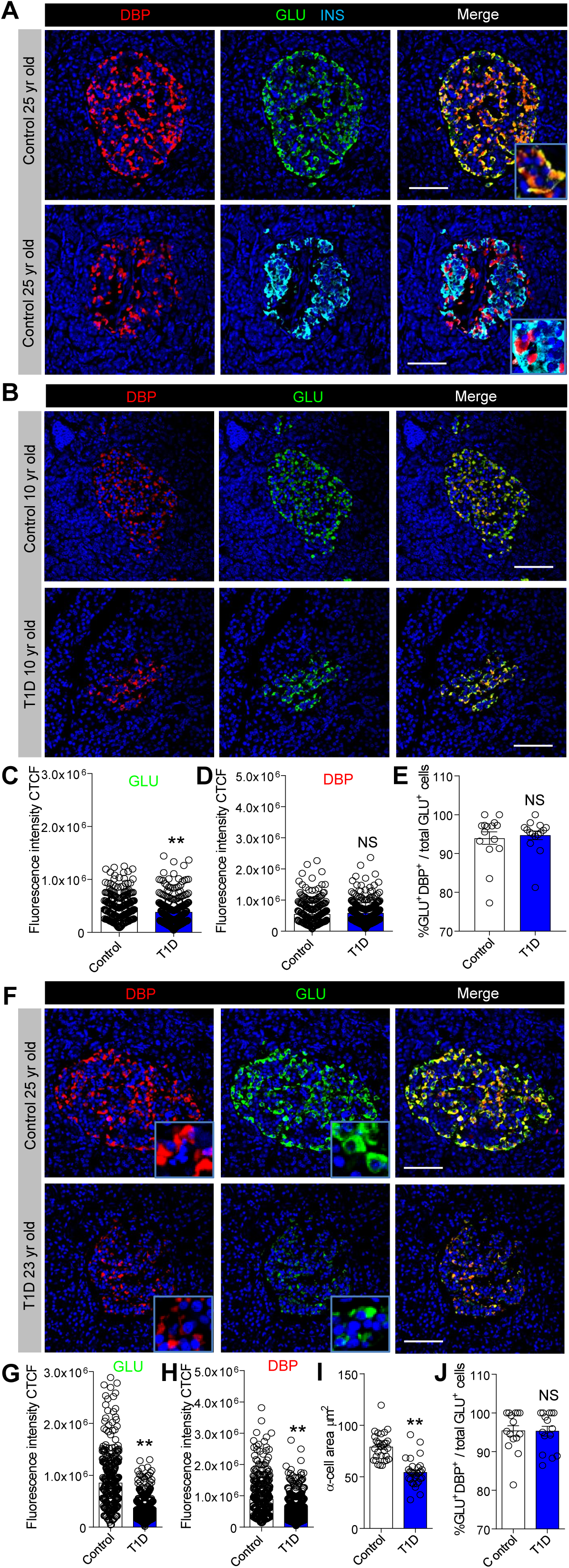
DBP expression is decreased in late-onset and longstanding type 1 diabetes. (A) Fluorescent immunohistochemistry showing strong expression of DBP in the α-cell compartment in human islets (inset shows a zoomed image) (n = 7 control donors). (B-E) Glucagon staining decreases slightly (B and C) in islets of donors with early-onset (<13 yrs old) T1D, but this is not associated with changes in DBP expression (B and D), or proportion of DBP+/GLU+ α-cells (E) (representative images are shown) (inset shows a zoomed image) (n = 300 cells, 30 islets, 3 T1D donors and age-matched controls; from the Exeter biobank) (Mann-Whitney test). (F-H) Glucagon (F and G) and DBP (F and H) expression are both decreased in islets of donors with late-onset (>13 yrs old) T1D (representative images are shown) (inset shows a zoomed image) (n = 400 cells, 40 islets, 4 T1D donors and age-matched controls) (Mann-Whitney test). (I and J) α-cell size (I), but not proportion of DBP+/GLU+ α-cells (J), is decreased in islets of donors with late-onset (>13 yrs old) T1D (n = 180 cells, 30 islets, 3 T1D donors and age-matched controls) (inset shows a zoomed image) (unpaired t-test). Scale bar = 42.5 µm. Bar graphs show scatter plot with mean ± SEM. **P<0.01 and NS, non-significant. DBP, Vitamin D-binding protein; GLU, glucagon; INS, insulin.

By contrast, glucagon levels were almost 2-fold lower in islets of late-onset T1D donors versus age-matched controls (Figure 5F and G), in line with previous reports of decreased α-cell mass during T1D (von Herrath et al., 2018). These changes were accompanied by a reduction in DBP expression (Figure 5H) and α-cell size (Figure 5I), although no differences were detected in the number of DBP+/GLU+ cells per islet (Figure 5J). We were able to confirm results in samples from IsletCore (Alberta), and also show that DBP levels consistently decrease in islets of donors with more longstanding T1D (Figure S3).

Immunohistochemistry of control islets showed that DBP expression increased with age, peaking at 18-32 years and remaining elevated thereafter (Figure 6A and B). Similar results were seen for glucagon expression (Figure 6C). Glucagon and DBP expression values for each individual donor are provided in Figure S4.

**Figure 6:**
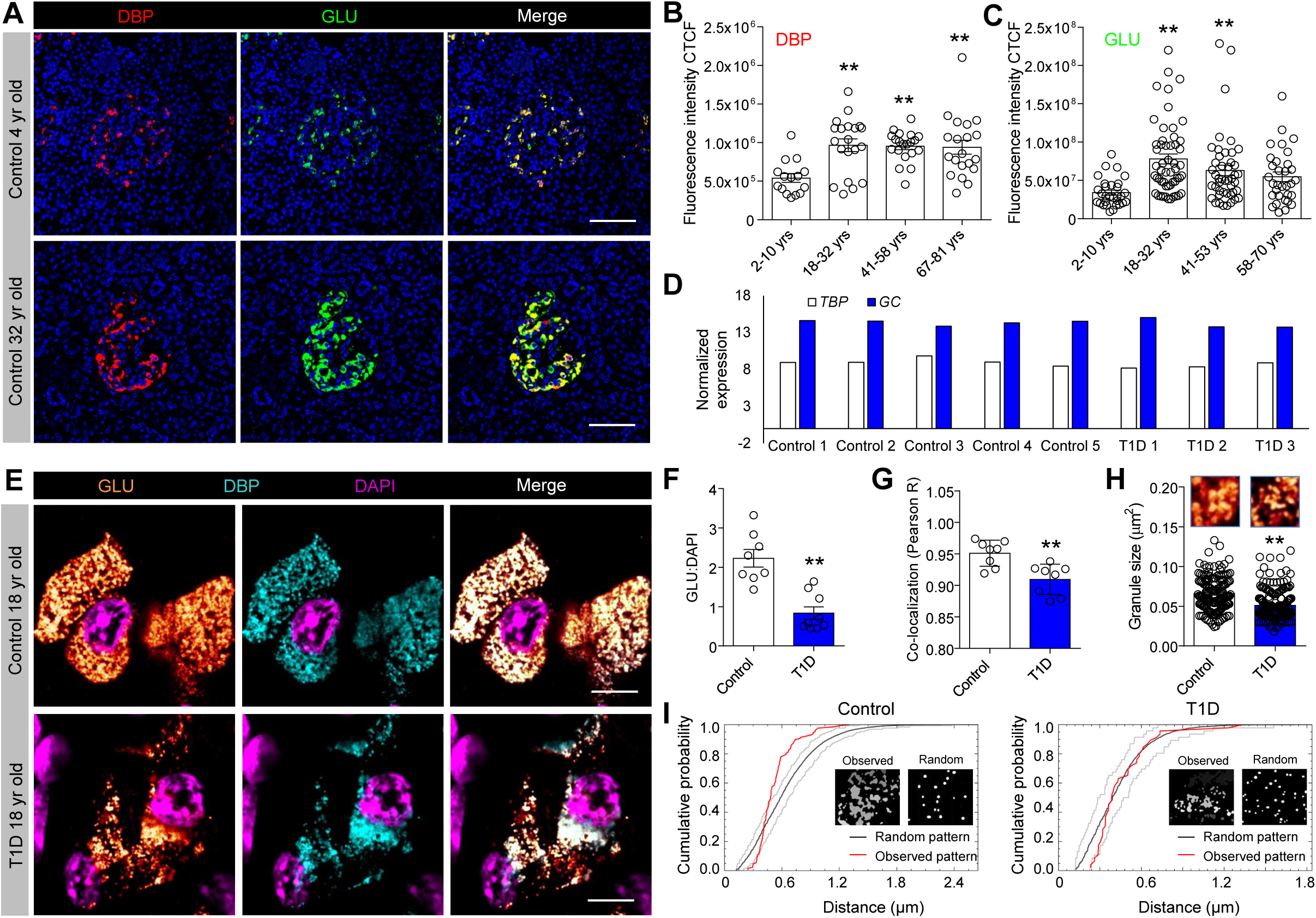
DBP expression increases with age and co-localizes with glucagon in granules. (A-C) DBP (A and B) and glucagon (A and C) expression increase with age in control donors (representative images are shown) (n = 15-53 islets per age group, 3 early-onset and 4 late-onset T1D donors together with age-matched controls) (one-way ANOVA with Tukey’s multiple comparison test). (D) Analysis of published RNA-seq datasets from purified α-cells (Brissova et al., 2018) shows no differences in transcript abundance for *GC* (encoding DBP) in control and T1D donors. Expression levels are normalized against *TBP*. Each individual donor is shown. Data were obtained from GEO: GSE106148. (E) Super-resolution images showing co-localization of DBP and glucagon within the same granule in α-cells of both control and late-onset T1D donors (representative images are shown) (n = 8 cells from 8 islets, 4 late-onset T1D donors together with age-matched controls). (F and G) The ratio of glucagon:DAPI (F) and DBP/glucagon co-localization strength (G) are lower in α-cells from donors with late-onset T1D (n = 8 cells, 4 late-onset T1D donors together with age-matched controls) (unpaired t-test). (H) Glucagon granule size is decreased in α-cells from donors with late onset T1D (magnified images from (E) are shown above each bar) (scale bar = 6 µm) (n = 160-200 granules from 4 islets, 4 late-onset T1D donors together with age-matched controls) (Mann-Whitney test). (I) G-function analysis on actual and simulated glucagon granule distribution showing a more random arrangement of glucagon granules in α-cells of T1D donors (actual and simulated distribution is inset). Scale bar = 42.5 µm. Bar graphs show scatter plot with mean ± SEM. **P<0.01 and NS, non-significant. DBP, Vitamin D-binding protein; GLU, glucagon; INS, insulin.; TBP, TATA-Box Binding Protein.

### Granular DBP content decreases in late-onset T1D

Analysis of a published RNA-seq dataset (Brissova et al., 2018) revealed no differences in *GC* expression in purified α-cells from control and T1D donors (20 to 53 years) (Figure 6D), suggesting that DBP might be post-transcriptionally regulated. As such, we investigated DBP and glucagon localization within human α-cells using super-resolution microscopy. Unexpectedly, DBP was found to be present in glucagon granules of control donors (Figure 6E), suggesting that DBP enters the secretory pathway and might function in an autocrine manner. Explaining the decrease in glucagon and DBP expression in late-onset T1D samples, a large reduction in the number of GLU+/DBP+ granules was detected in each α-cell (Figure 6F), accompanied by a small decrease in DBP/GLU co-localization (Figure 6G). Glucagon granules were also smaller and more randomly distributed in late-onset T1D α-cells (Figure 6H and I), suggestive of changes in the actin cytoskeleton, as demonstrated for mice.

## DISCUSSION

In the present study, we show that DBP is strongly expressed in murine and human α-cells. Loss of DBP leads to alterations in α-cell number and size, electrical activity and glucagon release. This is accompanied by changes in δ-cell mass, as well as alterations in β-cell function and insulin release. Linking these findings, DBP was found to decrease the availability of actin monomeric subunits for assembly into polymers. DBP expression levels were also decreased in islets of donors with late-onset or longstanding T1D, but not in children with early-onset disease.

Transcriptomic studies have consistently shown that *Gc* is highly-enriched in the mouse α-cell lineage (Adriaenssens et al., 2016; Cigliola et al., 2018; Qiu et al., 2017), similarly to data from humans (Ackermann et al., 2016; Segerstolpe et al., 2016). In keeping with these findings, our immunohistochemical analyses confirmed that DBP is predominantly expressed at the protein level in α-cells in both mouse and human. Careful inspection of images also detected faint DBP expression in β-cells, which was difficult to appreciate due to the high intensity of the DBP signal in α-cells. Notably, DBP expression was also absent in β-cells in DBP^-/-^ islets, and F-actin and G-actin were both altered across the islet. Thus, DBP is detectable in both α-cells and β-cells, with large differences in expression levels apparent between the two compartments. In addition, recent studies have shown that β-cells increase histone modification at the *Gc* locus in response to high fat feeding, with knockout of *Gc* protecting from β-cell dysfunction (Kuo et al., 2019). Thus, DBP protein expression in the β-cell compartment is low, but is likely to be upregulated under conditions of metabolic stress.

Intriguingly, Na^+^ currents, which contribute to action potential firing in α-cells, were increased in DBP^-/-^ α-cells. This was mirrored at the level of low glucose-stimulated Ca^2+^ rises, which were also increased by loss of DBP. Given that α-cell function was otherwise decreased across the board (i.e. defective glucagon secretion, activation by low glucose, electrophysiological identity, Na^+^ channel inactivation), increases in Na^+^ conductance and Ca^2+^ spiking amplitude are likely to reflect a maladaptive compensatory response.

Supporting the notion that F-actin is an important regulator of glucagon release (Hutchens and Piston, 2015), fiber size was increased in DBP^-/-^ animals, alongside decreases in monomeric G-actin content. Thus, DBP in the α-cell likely scavenges G-actin, preventing formation of F-actin polymers, which would otherwise suppress glucagon release. At the molecular level, F-actin has been shown to restrict basal insulin release (Kalwat and Thurmond, 2013), as well as maximal glucagon secretion, by acting as a physical constraint against granule exocytosis (Hutchens and Piston, 2015; Reissaus and Piston, 2017). Morphological evidence for this was provided in the current study by the observation that glucagon granules were distributed more diffusely in DBP^-/-^ islets. Furthermore, granule size was decreased, indicative of either preferential sequestration of immature granules, or effects of DBP on granule maturity. Cytoskeletal changes were also likely to be involved in the reduction in α-cell and δ-cell size, since assembly of actin filaments from monomers is critical for cell morphology (Pollard and Cooper, 2009).

β-cells displayed increased Ca^2+^ activity at low glucose in DBP^-/-^ islets. This however did not translate to elevated basal insulin release, probably because F-actin fiber thickness was also increased across β-cells, potentially acting as a barrier for unregulated granule exocytosis (Mziaut et al., 2016). While DBP levels were much lower in β-cells compared to α-cells, it could be argued that only small amounts are required to prevent actin polymerization given the high binding affinity for monomeric actin (Mc Leod et al., 1989). By contrast, the increase in glucose-stimulated insulin secretion seen in DBP^-/-^ islets is most likely due to lack of α-cell shut-off at high glucose, expected to positively reinforce β-cell function (Svendsen et al., 2018). Indeed, electrophysiological recordings showed that the Na^+^ channel inactivation switch tended to be impaired in DBP^-/-^ α-cells. An alternative mechanism might center on changes in glucose-stimulated actin remodelling in the β-cells themselves, for example by gelsolin (Tomas, 2006), which severs actin filaments but in competition with DBP.

Demonstrating the relevance of our studies for human disease, islets in pancreata from individuals with late-onset or longstanding T1D consistently showed decreased DBP expression, as well as a reduction in α-cell size. Super-resolution imaging showed that the majority of glucagon granules in human α-cells also contained DBP, with a sharp decrease in granular expression levels during T1D. A similar localization of DBP to secretory granules was reported in human neutrophils, together with release of the protein into the extracellular milieu (Kew et al., 1993). Since the *GC* transcriptional machinery is present in the α-cell, the source of this DBP is likely from de novo synthesis. It is also plausible that DBP is transported from the circulation into α-cells by megalin-mediated endocytosis, as reported in the kidney (Nykjaer et al., 1999), and that either this process, or liver production of DBP, is altered during T1D. However, it is difficult to envisage how endocytosis would lead to accumulation of DBP specifically in glucagon granules.

Interestingly, no changes in DBP expression were found in early-onset T1D donors. While the exact reasons for this are unknown, DBP was significantly lower in young versus old control donors without T1D. These data suggest that DBP expression and thus α-cell identity might not be fully specified until adolescence, meaning that DBP cannot be further downregulated in early-onset T1D. These data raise a number of interesting questions involving the known role of DBP as a novel autoantigen during T1D (Kodama et al., 2016). For example, does DBP only act as an auto-antigen in late-onset T1D patients? Is the decrease in DBP expression seen in late-onset T1D a consequence of auto-antigens, or another unrelated mechanism? If DBP is an auto-antigen in T1D, why do α-cells not die, or are the low DBP-expressing β-cells instead targeted? Could α-cells confer auto-immunity on β-cells through paracrine DBP signaling? How do these findings relate to polymorphic variants in *GC*, which are known to influence DBP action/levels, as well as 25(OH)D (Powe et al., 2013)? Further systematic studies in autoantigen-positive and -negative early and late-onset T1D donors, as well as individuals harboring GC risk alleles, will be required to address these questions. Nonetheless, our studies suggest that, together with adoption of a β-cell like transcriptional profile (Brissova et al., 2018), loss of DBP might contribute to the impaired glucagon secretion reported in T1D (Brissova et al., 2018; Marchetti et al., 2000).

We acknowledge a number of limitations in the present study. Firstly, the animals were globally deleted for DBP, which means that the effects of the protein specifically in α-cells could not be examined. However, DBP is highly expressed in α-cells, which validates our model. Moreover, use of DBP^-/-^ mice allowed us to uncover a hitherto underappreciated role of DBP in regulating β-cell and δ-cell function, which might also be important from a translational standpoint. In addition, a global deletion model would be more reflective of studies in humans bearing homozygous deletion of *GC* (Henderson et al., 2019). Secondly, although animals were fed a vitamin D-sufficient diet, we cannot completely exclude vitamin D-dependent effects of DBP. Suggesting that this is unlikely to be the case, a single individual harboring homozygous mutations in *GC* did not show symptoms consistent with vitamin D deficiency despite almost undetectable plasma 25(OH)D levels (Henderson et al., 2019). This argues for the free hormone hypothesis where DBP acts as a major vitamin D reservoir but only low levels are required for biological effects (Chun et al., 2014). Thirdly, morphometric analyses were based upon glucagon staining, which could lead to an underestimation of α-cell size in T1D samples, especially if the fewer detectable granules were not distributed evenly throughout the cytoplasm. Fourthly, although mice did not show a clear phenotype, further studies are warranted using in vivo models of α-cell stress, for example glucagon receptor antagonism to induce hyperplasia (Gu et al., 2009). Lastly, while a causal role for DBP in α-cell dysfunction is suggested by mouse studies, we cannot confidently assert the same in islets of human T1D donors where autoimmunity and species-differences come into play.

In summary, we show that DBP is critical for maintaining α-cell phenotype and glucagon secretion, with changes in expression apparent during late-onset and longstanding T1D. The stage is now set for investigating more widely how DBP influences islet function and disease risk in individuals with T1D and T2D.

## STAR METHODS

### KEY RESOURCES TABLE

#### CONTACT FOR REAGENT AND RESOURCE SHARING

Further information and requests for resources and reagents should be directed to and will be fulfilled by the Lead Contact, David J. Hodson.

#### STUDY DESIGN

No data were excluded unless the cells displayed a clear non-physiological state (i.e. impaired viability), and all individual data points are reported in the figures. The measurement unit is animal, batch of islets or donor, with experiments replicated independently at least three times. Samples and animals were allocated to treatment groups in a randomized manner to ensure that all states were represented in the different experiment arms. Investigators were blinded to animal and donor identity both during the experiment and subsequent analysis.

#### EXPERIMENTAL MODEL AND SUBJECT DETAILS

##### Mouse models

*DBP*^*-/-*^ mice were generated using a PGK-promoter/neomycin cassette to disrupt exon 5 of the mouse *Gc* gene, as described (Safadi et al., 1999). We used these animals rather than the KOMP repository strain (Gctm1^(KOMP)Vlcg^; VG11723), since they have been subjected to thorough phenotypic validation and show loss of serum DBP protein, as well as 25(OH)[^3^H]D_3_ binding (Safadi et al., 1999). Animals were maintained in a specific-pathogen free facility with ad lib access to regular chow (which contains 1000 U/kg cholecalciferol) and water. All studies were performed with 6-15 week-old male and female animals, and regulated by the Animals (Scientific Procedures) Act 1986 of the U.K. Approval was granted by the University of Birmingham’s Animal Welfare and Ethical Review Body.

##### Human donors

Formalin-fixed paraffin-embedded pancreata were obtained from the Exeter Archival Diabetes Biobank (http://foulis.vub.ac.be/) or the Alberta Diabetes Institute IsletCore (quality control and phenotyping data is available for each preparation (via www.isletcore.ca and iidp.coh.org). Studies with human tissue were approved by the University of Birmingham Ethics Committee, as well as the National Research Ethics Committee (REC reference 16/NE/0107, Newcastle and North Tyneside, U.K.). Donor characteristics are reported in Table S1.

## METHOD DETAILS

### Islet isolation and culture

Animals were euthanized by a cervical dislocation, before isolation of islets using collagenase digestion (1 mg/ml, SERVA NB8; amsbio Cat# 17456.01) and Histopaque or Ficoll-Paque gradient separation. Islets were maintained at 37°C and 5% CO_2_ in RPMI medium containing 10% FCS, 100 units/mL penicillin, and 100 μg/mL streptomycin.

### Gene expression

Relative mRNA abundance was determined using an Applied Biosystems 7500 or 7900HT instrument and PowerUp SYBR Green Master Mix (Thermo Fisher Scientific Cat# A25742). Fold-change mRNA expression was calculated versus *Ppia* by using the 2^−ΔΔCt^ method. For primer sequences, see the Resource Table.

### Glucagon and insulin assays

Batches of 10 islets were pre-incubated in either 10 mM or 3 mM glucose for 1 hour at 37°C in buffer containing (in mmol/L) 120 NaCl, 4.8 KCl, 24 NaHCO_3_, 0.5 Na_2_HPO_4_, 5 HEPES, 2.5 CaCl_2_, 1.2 MgCl_2_, and 3–17 D-glucose + 0.1% BSA. For glucagon secretion, islets were incubated in 10 mM, 0.5 mM or 0.5 mM glucose + 5 μm epinephrine for 1 hour at 37°C. Glucagon and insulin secreted into the supernatant was measured using specific ultrasensitive HTRF assay (glucagon; Cisbio Cat# 62CGLPEG) (insulin; Cisbio Cat# 62IN2PEG). Total glucagon was measured from islets lysed in acid ethanol. Insulin was measured similarly, but using batches of 10 islets sequentially incubated in 3 mM glucose, 17 mM glucose and 17 mM glucose + 10 mM KCl for 30 minutes at 37°C. In all cases, values are normalized against total glucagon/insulin for each individual experiment to account for differences in α-cell/β-cell proportion with treatment and islet size (Henquin, 2019).

### Immunostaining of mouse tissue

Pancreata were fixed in 10% formalin overnight, before dehydration and wax embedding. Sections were blocked with PBS-T + 1% BSA for 1 hour and incubated with primary antibodies overnight at 4°C. Following washing in PBS-T + 0.1% BSA, secondary antibodies were applied for 2 hours at room temperature. Primary antibodies were rabbit anti-insulin 1:500 (Cell Signaling Technology Cat# 3014, RRID:AB_2126503), guinea pig anti-insulin 1:50 (Abcam Cat# ab7842, RRID:AB_306130), mouse monoclonal anti-glucagon 1:2000 (Sigma-Aldrich Cat# G2654, RRID:AB_259852), rabbit anti-glucagon 1:100 (Sigma-Aldrich Cat# SAB4501137, RRID:AB_10761583), mouse anti-somatostatin 1:1000 (Thermo Fisher Scientific Cat#14-9751-80, RRID:AB_2572981), rabbit anti-DBP 1:1000 (Sigma-Aldrich Cat# HPA019855, RRID:AB_1849545), guinea pig anti-PDX1 1:200 (Abcam Cat# ab47308, RRID:AB_777178), rabbit anti-MafA 1:1000 (Bethyl laboratories Cat# IHC-00352, RRID:AB_1279486), and mouse anti-PCNA 1:500 (Cell Signaling Technology Cat# 2586, RRID:AB_2160343). We note that the rabbit anti-DBP antibody (Sigma-Aldrich Cat# HPA019855, RRID:AB_1849545) was developed and validated by the Human Protein Atlas project, passing multiple quality controls (https://www.proteinatlas.org/ENSG00000145321-GC/antibody#protein_array).

Secondary antibodies were goat anti-rabbit Alexa Fluor 633 (Thermo Fisher Scientific Cat# A-21052, RRID:AB_2535719), goat anti-rabbit Alexa Fluor 488 (Thermo Fisher Scientific Cat# R37116, RRID:AB_2556544), goat anti-guinea pig Alexa Fluor 488 (Thermo Fisher Scientific Cat# A-11073, RRID:AB_2534117), goat anti-mouse Alexa Fluor 488 (Thermo Fisher Scientific Cat# A-11029, RRID:AB_138404), goat anti-guinea pig Alexa Fluor 568 (Thermo Fisher Scientific Cat# A-11075, RRID:AB_2534119) 1:1000. Fixed islets were incubated with Phalloidin-488 (Abcam Cat# ab176753) and DNAseI-594 (Invitrogen Cat# D12372) for 2 hours at room temperature to stain F-actin and G-actin.

Images were captured using either Zeiss LSM780 or LSM880 meta-confocal microscopes, the latter equipped with an Airyscan super-resolution module. Excitation was delivered at λ = 488 nm, λ = 568 and λ = 633 nm for Alexa Fluor 488, Alexa Fluor 568 and Alexa Fluor 633, respectively. Emitted signals were detected using a GaAsP PMT detector at λ = 498–559, nm λ = 568–629 and λ = 633–735 nm for Alexa Fluor 488, Alexa Fluor 568 and Alexa Fluor 633, respectively. Super-resolution images were subjected to online deconvolution processing using Zen Black (Zeiss Microscopy).

### Immunostaining of human tissue

Tissue was obtained from individuals with T1D and their age-matched controls. Donor details are provided in Table S1. Samples were dewaxed and rehydrated before antigen retrieval and blocking with 5% normal goat serum. Primary antibodies were guinea pig anti-insulin 1:700 (Agilent Cat# A0564, RRID:AB_10013624), mouse anti-glucagon 1:2000 (Abcam Cat# ab10988, RRID:AB_297642) or mouse monoclonal anti-glucagon 1:2000 (Sigma-Aldrich Cat# G2654, RRID:AB_259852), and rabbit anti-DBP 1:500 (Sigma-Aldrich Cat# HPA019855, RRID:AB_1849545). Secondary antibodies were goat anti-guinea pig Alexa Fluor 647 (Thermo Fisher Scientific Cat# A-21450, RRID:AB_2735091), goat anti-mouse Alexa Fluor 555 (Thermo Fisher Scientific Cat# A-11075, RRID:AB_2534119), and goat anti-rabbit Alexa Fluor 488 at 1:400 (Thermo Fisher Scientific Cat# R37116, RRID:AB_2556544).

Images were captured using Zeiss LSM780 and LSM880 meta-confocal microscopes, as above. Excitation was delivered at λ = 488 nm, λ = 568 nm and λ = 633 nm for Alexa Fluor 488, Alexa Fluor 555 and Alexa Fluor 647 nm, respectively. Emitted signals were detected using a GaAsP PMT detector at λ = 498–561 nm, λ = 564–617 nm and λ = 641–691 nm for Alexa Fluor 488, Alexa Fluor 555 and Alexa Fluor 647 nm, respectively.

### Analysis of α-cell and β-cell mass

Pancreatic sections for determination of α-cell and β-cell mass were stained as above, before scanning and digitization using a Zeiss Axio Scan.Z1. Excitation was delivered at λ = 453-485 nm and λ = 590-650 nm for Alexa Fluor 488 and Alexa Fluor 647, respectively. Emitted signals were detected using an Orca Flash 4.0 at λ = 507-546 nm and λ = 663-738 nm for Alexa Fluor 488 and Alexa Fluor 647, respectively. Overall 408 separate images were captured for each pancreas section using a 20 x / 0.8 NA objective, before compilation into a single image using Zen lite 2012.

### Ca^2+^ imaging

Islets were loaded with Fura2 (HelloBio HB0780-1mg) before imaging using a Crest X-Light spinning disk system coupled to a Nikon Ti-E base and 10 x / 0.4 air objective. Excitation was delivered at λ = 340 nm and λ = 385 nm using a FuraLED system, with emitted signals detected at λ = 470–550 nm. Traces were presented as the emission ratio at 340 nm and 385 nm (i.e. 340/385). HEPES-bicarbonate buffer was used, containing (in mmol/L) 120 NaCl, 4.8 KCl, 24 NaHCO_3_, 0.5 Na_2_HPO_4_, 5 HEPES, 2.5 CaCl_2_, 1.2 MgCl_2_, and 0.5–17 D-glucose.

### Electrophysiology

Whole-cell currents were recorded in intact islets using the standard whole-cell configuration, as previously described (Briant et al., 2018). Measurements were performed using an EPC-10 patch-clamp amplifier and Patchmaster software (HEKA Electronics). Currents were filtered at 2.9 kHz and digitized at more than 10 kHz. Currents were compensated for capacitive transients and leak current subtraction was conducted. The extracellular solution consisted of (in mM) 138 NaCl, 5.6 KCl, 1.2 MgCl_2_, 5 HEPES (pH 7.4 with NaOH), 2.6 CaCl_2_ and 1 D-glucose. The intracellular solution contained (mM) 125 KCl, 1 CaCl_2_, 1 MgCl_2_, 5 HEPES, 3 MgATP and 10 EGTA (KOH buffered). All chemicals were from Sigma-Aldrich UK. Recordings with an access resistance of < 50 mΩ were used for analysis in MATLAB. The logistic regression model identifying cell type was implemented in MATLAB as previously described (Briant et al., 2017a).

### QUANTIFICATION AND STATISTICAL ANALYSIS

#### Correlation analysis

Correlation analysis was performed using matrix binarization analyses (Hodson et al., 2012). Intensity over time traces were smoothed using Hilbert-Huang empirical mode decomposition, and a 20% threshold used to binarize cells according to activity status. Co-activity was assessed using the equation C_ij_ = T_ij_/√T_i_T_j_ where C is a correlation coefficient, T_i_ and T_j_ is the period spent ON for each cell, and T_ij_ is the period both cells spend on together. Significance (P<0.01) was calculated versus the randomized dataset using a permutation step based. Highly connected cells were identified based on their position (60-100% range) within a probability-distribution, before construction of functional connectivity using link number and Euclidean coordinates.

#### Image analysis

F-actin, G-actin and DBP expression levels were analyzed using integrated density (area x mean fluorescence intensity), which accounts for the influence of cell size on fluorophore emission intensity for a given pixel (i.e. intensity of *n* fluorescent molecules will increase as a function of area^-1^). Corrected total cell fluorescence (CTCF) was then calculated as follows: integrated density – (area of selected cell x mean background fluorescence) (Gavet and Pines, 2010). Quantification of α-cell, β-cell and δ-cell area and number was performed on binarized images using ImageJ (NIH) and Threshold, Nucleus Counter and Particle Analysis plugins.

Glucagon granule distribution was analyzed using the G-function, which measures the distance from any position to the nearest object of interest compared to a random distribution of the same measured objects (FIJI Spatial Statistic 2D/3D plugin) (Zimmer et al., 2010). A left shift away from the mean +/-95% confidence intervals indicates a less random or more clustered organization.

Linear adjustments to brightness and contrast were applied to representative images, with intensity values maintained between samples to allow accurate cross-comparison. For super-resolution images, the following FIJI look-up-tables were used: NanoJ-Orange, cyan and magenta.

#### Statistical analysis

Data normality was assessed using D’Agostino-Person test. Unpaired/paired Students t-test or Mann-Whitney test were used for pairwise comparisons. Multiple interactions were determined using one-way or two-way ANOVA followed by Tukey’s, Dunnett’s, Bonferonni’s or Sidak’s post-hoc tests (accounting for degrees of freedom). Analyses were conducted using GraphPad Prism or Excel software

## Data availability

The datasets generated and/or analyzed during the current study are available from the corresponding author upon reasonable request.

## ACKNOWLEDGMENTS

D.J.H. was supported by MRC (MR/N00275X/1 and MR/S025618/1) and Diabetes UK (17/0005681) Project Grants. This project has received funding from the European Research Council (ERC) under the European Union’s Horizon 2020 research and innovation programme (Starting Grant 715884 to D.J.H.). L.J.B.B. was supported by a Sir Henry Wellcome Postdoctoral Fellowship (Wellcome Trust, 201325/Z/16/Z) and a Junior Research Fellowship from Trinity College, Oxford. P.E.M. was funded by a Foundation Grant from the Canadian Institutes of Health Research (grant 148451). N.G.M. and S.J.R. were supported by Diabetes UK (15/0005156 and 16/0005480), MRC (MR/P010695/1) and JDRF (2-SRA-2018-474-S-B) Project Grants. The authors thank the Human Organ Procurement and Exchange and Trillium Gift of Life Network programs for their efforts in procuring pancreases for research at the Alberta Diabetes Institute.

## AUTHOR CONTRIBUTIONS

K.V., D.N., D.L., N.H.F., F.A., M.J.R. and G.dS.X. performed in vitro experiments and analyzed data. K.V., S.H., D.L. G.G.L., performed in vivo studies and analyzed data. L.J.B.B. performed electrophysiological studies and analyzed data. J.E.M.F. and P.E.M. provided human pancreas samples. I.A. performed bioinformatic analysis. K.V., D.J.H., C.F., N.G.M. and S.J.R. performed immunohistochemical analysis of human samples.. M.H. and D.J.H. conceived, designed and supervised the studies. K.V., M.H. and D.J.H. wrote the paper with input from all authors.

## DATA AVAILABILITY

All data necessary to understand, assess and extend the conclusions of the manuscript are available upon reasonable request.

