## Supplementary Material for "Vitamin D-binding protein is required for the maintenance of α-cell function and glucagon secretion"

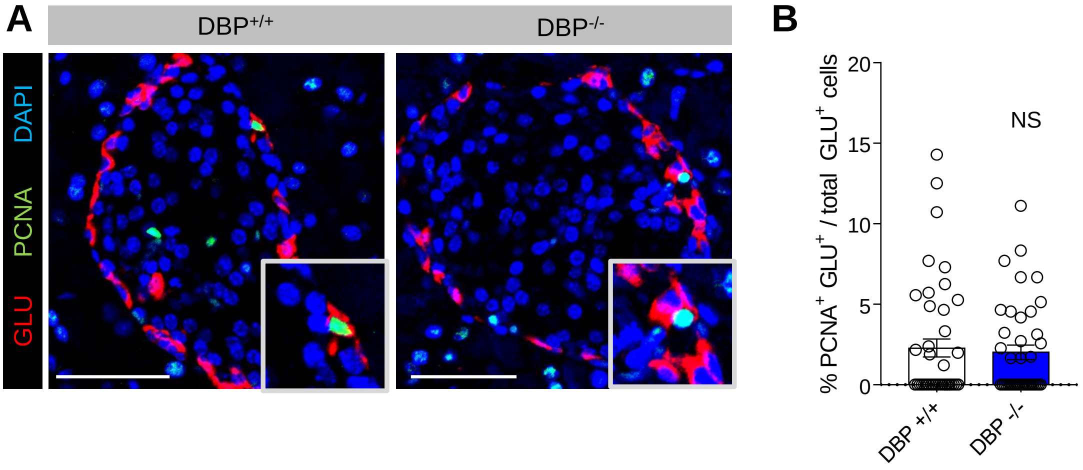


**Supplemental Figure 1 (related to Figure 2): No changes in α-cell turnover are detected following loss of DBP**

(A) Representative images showing PCNA and glucagon (GCG) staining in in islets of $DBP^{+/+}$ and $DBP^{-/-}$ mice (n = 4 animals).

(B) The number of PCNA-positive α-cells is similar in islets of $DBP^{+/+}$ and $DBP^{-/-}$ mice (n = 4 animals)

Bar graph shows scatter plot with mean ± SEM. NS, non-significant. DBP, Vitamin D-binding protein; GLU, glucagon; PCNA, proliferating cell nuclear antigen.

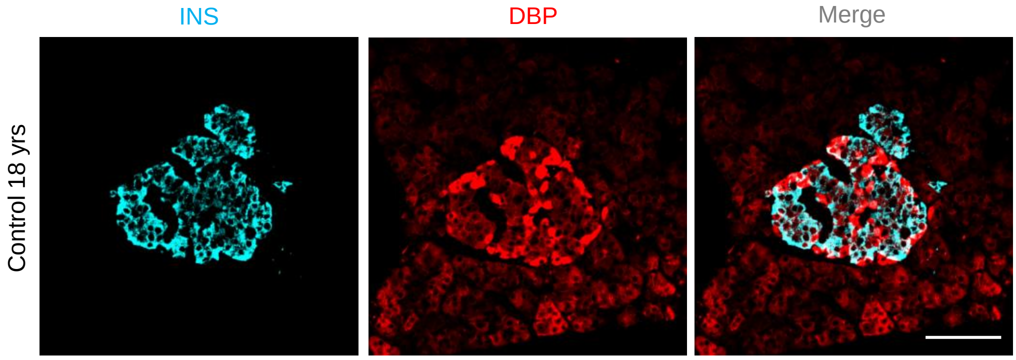


**Supplemental Figure 2 (related to Figure 5): DBP is present at low levels in β-cells**

DBP can be detected in insulin-positive cells, as well as exocrine tissue, in control human donors. Due to the strength of DBP expression in α-cells, the image has been overexposed to allow visualization of DBP in the glucagon-negative compartment (scale bar = 42.5 µm ) (representative images from n = 3 control donors). DBP, Vitamin D-binding protein; GLU, glucagon; INS, insulin.

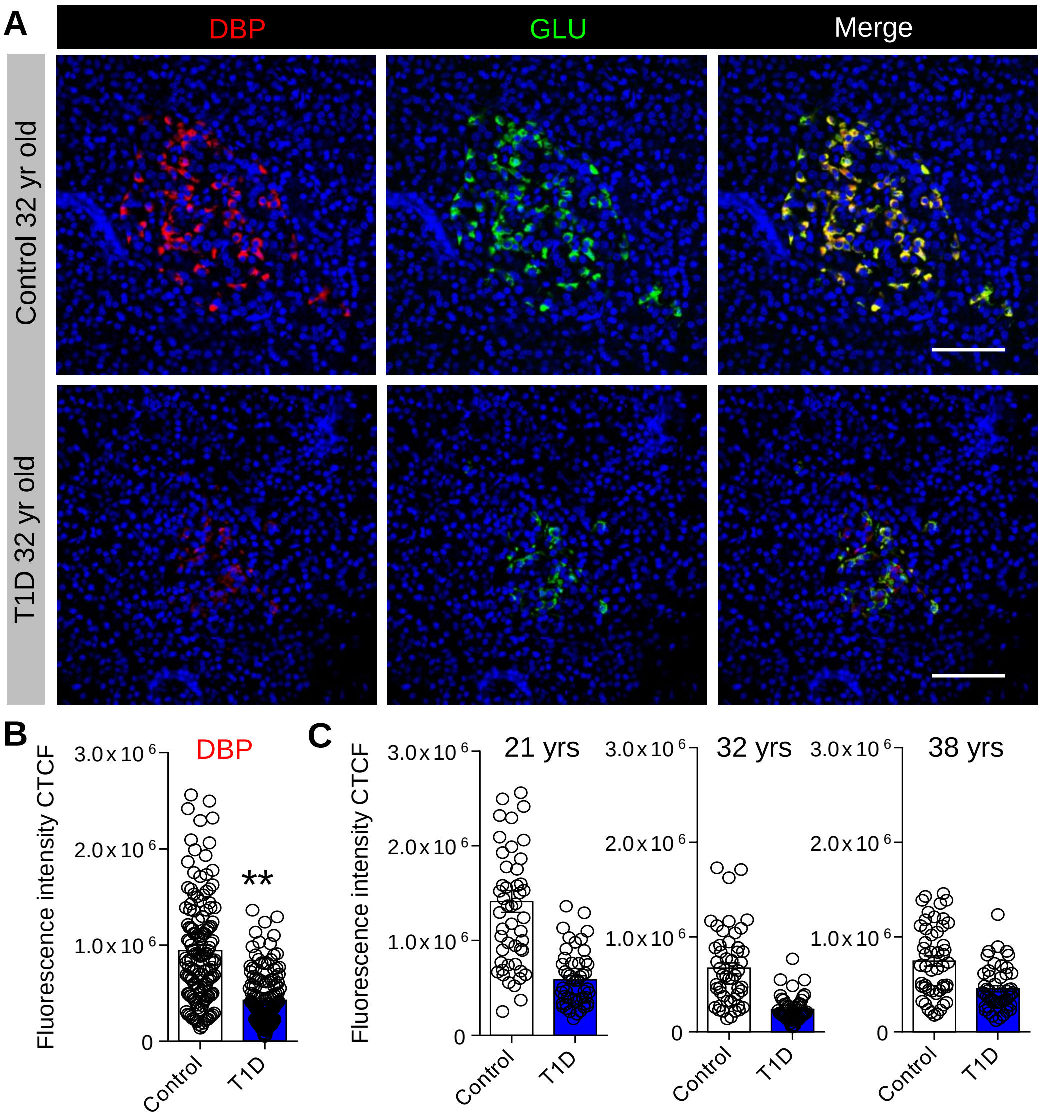


**Supplemental Figure 3 (related to Figure 5): DBP is decreases in islets of donors with more longstanding T1D.**

(A) Representative images showing DBP expression in islets of donors with more longstanding T1D, together with their age-matched controls, both obtained from the IsletCore (Alberta) biobank (3 T1D donors and 3 age-matched controls).

(B and C) DBP levels are decreased in adult donors with longstanding T1D, as shown by mean (B) and individual donor (C) data (n = 150 cells, 30 islets, 3 T1D donors and 3 age-matched controls) (unpaired t-test).

Scale bar = 42.5 µm. Bar graphs show scatter plot with mean ± SEM. **P<0.01 and NS, non-significant. DBP, Vitamin D-binding protein; GLU, glucagon; INS, insulin.

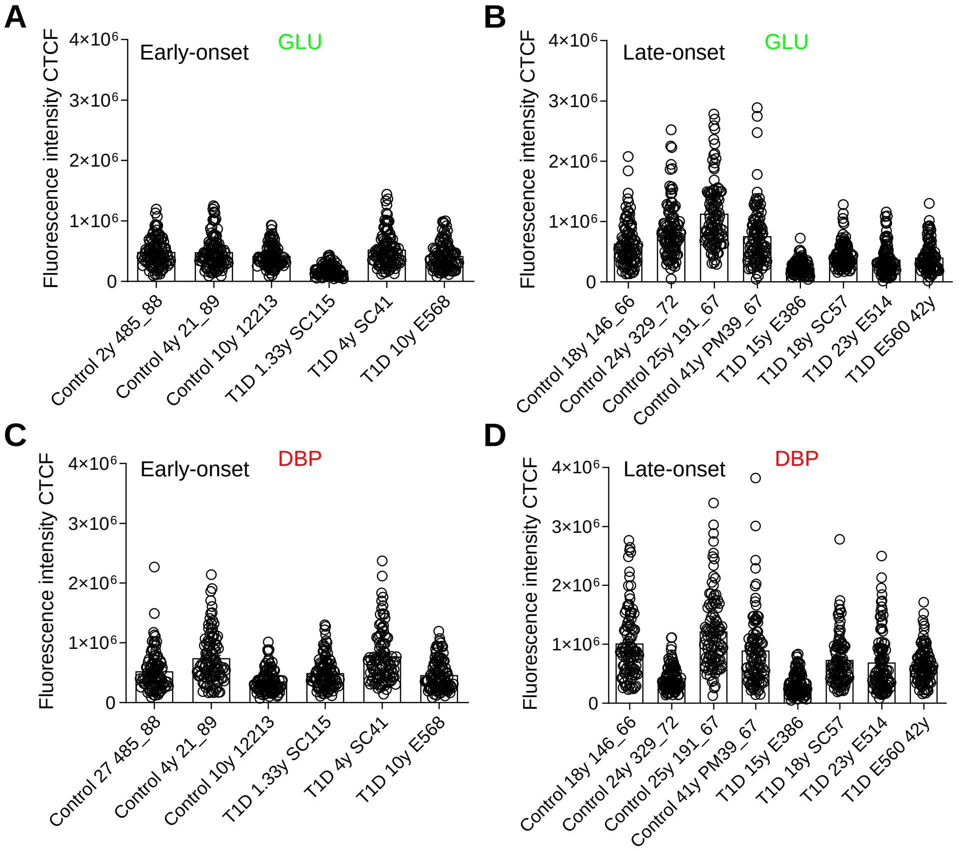


**Supplemental Figure 4 (related to Figure 5): Individual donor glucagon and DBP expression levels**

(A and B) Expression levels of glucagon in individual early-onset (A) and late-onset (B) T1D donors together with their age-matched controls (n = 100 cells, 10 islets from each donor).

(C and D) Expression levels of DBP in individual early-onset (C) and late-onset (D) T1D donors together with their age-matched controls (n = 100 cells, 10 islets from each donor).

Bar graphs show scatter plot with mean ± SEM. DBP, Vitamin D-binding protein; GLU, glucagon.

| Tissue | Donor case ID | Age (years) | Sex | Cohort | Duration |
|---|---|---|---|---|---|
| Control | 485/88 | 2 | F | Exeter | |
| Control | 21/89 | 4 | F | Exeter | |
| Control | 12213 | 10 | unknown | Exeter | |
| Control | 146/66 | 18 | unknown | Exeter | |
| Control | 329/72 | 24 | unknown | Exeter | |
| Control | 191_67 | 25 | M | Exeter | |
| Control | 447/71 | 32 | F | Exeter | |
| Control | PM39/67 | 41 | unknown | Exeter | |
| Control | 330_71 | 47 | M | Exeter | |
| Control | 3730A/88 | 53 | F | Exeter | |
| Control | 9310/08 | 58 | F | Exeter | |
| Control | 224/66 | 67 | M | Exeter | |
| Control | 36/66 | 70 | F | Exeter | |
| Control | 424/67 | 70 | M | Exeter | |
| Control | R187 | 21 | M | Alberta | |
| Control | R294 | 32 | M | Alberta | |
| Control | R044 | 38 | M | Alberta | |
| T1D | SC115 | 1.33 | F | Exeter | 3 days |
| T1D | SC41 | 4 | F | Exeter | 3 wks |
| T1D | E568 | 10 | M | Exeter | 2 days |
| T1D | E386 | 15 | M | Exeter | 6 mths |
| T1D | SC57 | 18 | F | Exeter | < 1 wk |
| T1D | E514 | 23 | M | Exeter | 2 wks |
| T1D | E560 | 42 | n/a | Exeter | n/a |
| T1D | R015 | 21 | M | Alberta | n/a |
| T1D | R079 | 32 | M | Alberta | 17 yrs |
| T1D | R035 | 38 | F | Alberta | 15 yrs |

**Supplemental Table 1:** List of control and T1D donors used for immunohistochemical analyses in Figure 5 & 6. M, male; F, female; n/a, not available.
